# Side-binding proteins modulate actin filament dynamics

**DOI:** 10.1101/008128

**Authors:** Alvaro H. Crevenna, Marcelino Arciniega, Aurélie Dupont, Kaja Kowalska, Oliver F. Lange, Roland Wedlich-Söldner, Don C. Lamb

**Affiliations:** Physical Chemistry, Department for Chemistry and Biochemistry and Center for Nanoscience (CeNS), Ludwig-Maximilians-Universität München, Butenandtstraße 5-13, Haus E, 81377, München, Germany; Cellular Dynamics and Cell Patterning, Max Planck Institute of Biochemistry, Am Klopferspitz 18, 82152, Martinsried, Germany; Max Planck Institute of Biochemistry, Am Klopferspitz 18, 82152, Martinsried, Germany; Department Chemie, Technische Universität München, Lichtenbergstraße 4, 85748 Garching, Germany; Biomolecular NMR and Munich Center for Integrated Protein Science, Technische Universität München, Lichtenbergstraße 4, 85748 Garching, Germany; Munich Center for Integrated Protein Science (CiPSM), Ludwig-Maximilians-Universität München, Butenandtstraße 11, 81377 München, Germany; Institute of Structural Biology, Helmholtz Zentrum München, 85764 Neuherberg, Germany; Department of Physics, University of Illinois at Urbana-Champaign, 1110 West Green Street, Urbana, IL 61801, USA; Present address: Institute of Cell Dynamics and Imaging, Von-Esmarch-Straße 56, 48149, Münster, Germany

## Abstract

Actin filament dynamics govern many key physiological processes from cell motility to tissue morphogenesis. A central feature of actin dynamics is the capacity of the filament to polymerize and depolymerize at its ends in response to cellular conditions. It is currently thought that filament kinetics can be described by a single rate constant for each end. Here, using direct visualization of single actin filament elongation, we show that actin polymerization kinetics at both filament ends are strongly influenced by proteins that bind to the lateral filament surface. We also show that the less dynamic end, called the pointed-end, has a non-elongating state that dominates the observed filament kinetic asymmetry. Estimates of filament flexibility and Brownian dynamics simulations suggest that the observed kinetic diversity arises from structural alteration. Tuning filament kinetics by exploiting the natural malleability of the actin filament structure may be a ubiquitous mechanism to generate the rich variety of observed cellular actin dynamics.

## Introduction

Central cellular processes such as cell migration, cytokinesis, endocytosis and mechanosensation depend critically on actin-based force generation and actin filament turnover (Lecuit et al., 2011; Pollard and Borisy, 2003). The molecular basis of actin filament turnover derives from the association and dissociation of monomers from each filament end and depends on the nucleotide (ATP, ADP · Pi or ADP) bound to the actin monomer (Pollard, 1986). The filament is kinetically asymmetric, where one end (called the barbed-end) is observed to grow an order of magnitude faster than the other end (the pointed-end) (Pollard, 1986). In addition, the critical concentration for polymerization is different for the two ends. The origin of the asymmetry is not fully understood. Measurements of filament elongation as a function of solution viscosity (Drenckhahn and Pollard, 1986) and particle-analysis from cryo-electron microscopy (Narita et al., 2011) suggest the existence of a non-elongating state at the pointed-end. However, direct evidence for a non-elongating state via imaging of filament elongation using total internal reflection fluorescence ‘TIRF’ microscopy has not been observed (Fujiwara et al., 2007; Kuhn and Pollard, 2005). The dynamics of the pointed end plays an important role in both the origin of the differences in critical concentration at the two ends in the presence of ATP as observed (Fujiwara et al., 2007; Pollard, 1986); and in filament treadmilling, where, barbed-end growth and pointed-end shrinking occur simultaneously (Bugyi and Carlier, 2010). Thus, we have focused on performing an accurate and detailed analysis of both barbed-end and pointed-end dynamics using TIRF microscopy.

In cells, a large number of proteins interact with actin filaments, either at the ends or with the lattice. End-binding proteins, generally referred to as capping proteins, regulate actin dynamics by limiting elongation (at the barbed-end) or serving as anchor points (for the pointed-end). Side-binding proteins, on the other hand, are much more diverse encompassing myosin motors, cross-linkers or bundlers as well as severing proteins. The interaction of the actin filament with a particular subset of proteins defines the molecular composition, architecture and overall turnover of sub-cellular arrays such as stress-fibers and filopodia. Some of these arrays are tightly packed (Jasnin et al., 2013) and dynamics of the filaments may be influenced by the local environment. The mechanisms of how some proteins are recruited to these structures while others are excluded are the subject of intense research (Cai et al., 2008; Hansen et al., 2013). Although the overall filament dynamics have been thought to be sensitive to the concentration of the side-binding protein (Breitsprecher et al., 2009a), it is not understood how and to what extent side-binding proteins alter filament kinetics, structure and flexibility.

Here, we used total internal reflection fluorescence ‘TIRF’ microscopy to study the effect of side-binding proteins on the dynamics of actin filament growth *in vitro*. We chose three cross-linking proteins and one motor protein to represent the large variety of interacting proteins and used them to tether filaments directly to the surface of a glass slide for visualization. We used the chemically inactivated myosin II motor protein (NEM-myosin) as it is the standard choice for this type of assay (Kuhn and Pollard, 2005). The filamin protein (Kueh et al., 2008) was used, which is an important player in cellular mechanosensing that is evolutionary-conserved (Razinia et al., 2012), as its use as a tether has recently generated some debate (Mullins, 2012; Niedermayer et al., 2012). Additionally, we selected: α-actinin, a molecule that, together with myosin II, forms stress fibers (Langanger et al., 1986); and VASP, a protein that localizes to areas of dynamic actin reorganization such as filopodia and the lamelipodium (Rottner et al., 1999). By carrying out these assays with several proteins that bind to the side of actin filaments, we were able to explore the possible range of modulation available to actin filament dynamics and delineate intrinsic filament properties.

## Results

### Kinetic modulation at the barbed-end

Fluorescently labeled actin was used to visualize the growth of actin filaments (Figure 1a-b) using TIRF microscopy. In this technique, single actin filaments are tethered to a glass surface via a side-binding protein and their growth and/or shrinkage is monitored in real time (Figure 1a-b). From each frame, the filament is extracted into a kymograph. The position of each end of the filament was then determined by fitting an error function (Demchouk et al., 2011) to each line of the kymograph (For details, see Supporting Information and Figure S1). This end-detection method provides a more accurate determination of the filament length and thereby a more reliable estimate of the instantaneous elongation velocity compared to methodologies used previously (Figure S1).

**Figure 1.**
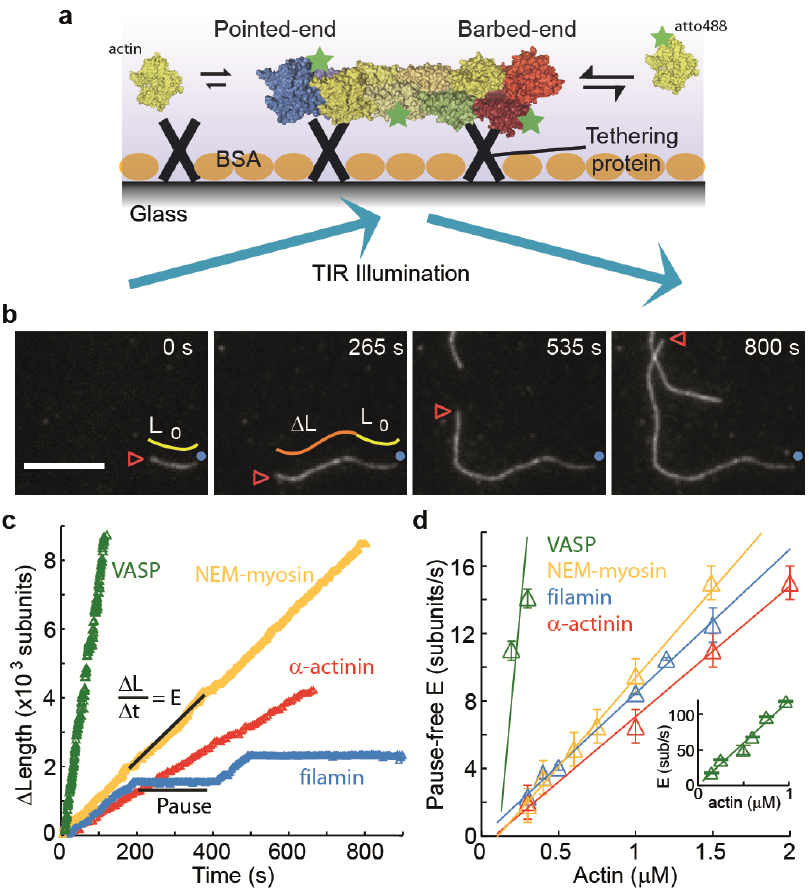
The dependence of the barbed-end kinetics on the side-binding protein. **a**, Schematic of total internal reflection (TIR) illumination and single actin-filament imaging tethered to a glass surface. Filaments grow from the addition of subunits at either the barbed-or the pointed-end. **b**, Selected frames from a movie showing the growth of single actin filament that is tethered to the surface via α-actinin. The barbed-end is marked by a red arrowhead and the pointed-end by a blue dot. The elapsed time is given in seconds. Scale bar: 5 μm. *L*_0_ and Δ*L* are the initial filament length and the change in length respectively. **c**, Δ*L* as a function of time for single filaments grown on surfaces with different tethering proteins. **d**, Elongation velocity (E) as a function of actin concentration in solution for different tethering proteins (Inset, zoom out to display the VASP values). The elongation velocity was determined from the slope of the graphs of Δ*L* versus time in regions where no pauses were observable. Error bars represent s.e.m. (n >20). Tether density here is ∼2,000 molecules/μm^2^.

The single-filament elongation experiments showed that the barbed-end grew at a constant velocity with occasional pauses for all constructs measured while the barbed-end elongation velocity varied depending on the particular side-binding protein used (Figure 1c, d). The elongation velocity ‘*E*’ at the barbed-end was the fastest with VASP and the slowest with α-actinin (Figure 1c). By varying the free actin concentration from 0.3-2 μM, we estimated the barbed-end association and dissociation rates, *k*_on_ and *k*_off_ respectively (Figure 1d), using only the periods of elongation (i.e. E > 1.5 sub·s^−1^, referred hereafter to as ‘kinetically active’ phases). Compared to the previously reported value (Pollard, 1986) of 11.6 sub·μM^−1^·s^−1^ for actin only in the absence of tethering proteins, we found a higher value of *k*_on_ in the presence of VASP and a lower value when NEM-myosin, α-actinin or filamin were used (Figure 1d). Extrapolating the elongation velocity as a function of actin concentration to zero actin provides an estimate of the dissociation rate, *k*_off_, of ATP-actin at the barbed-end (Table 1). In the presence of filamin, *k*_off_ is indistinguishable from zero whereas, in the presence of VASP, *k*_off_ increased compared to that value in the presence of NEM-myosin. The estimated *k*_off_ we measured in the presence of NEM-myosin (1.6 ± 0.5 s^−1^) was in agreement with the previously reported value of 1.4 s^−1^ (Pollard, 1986) whereas in the presence of α-actinin, was lower than 1.4 s^−1^. The ratio of inferred dissociation rates to the calculated association rate (i.e. *k*_off_/*k*_on_) is the critical concentration at which polymerization will occur and has been estimated to be ∼150 nM for the barbed-end (Pollard, 1986). We find a similar value (∼0.2 μM) for filaments elongated in the presence of VASP, α-actinin and NEM-myosin, but close to zero for filamin. These results demonstrate that ATP-actin kinetics at the barbed-end is sensitive to the particular side-binding protein interacting with the filament.

**Table 1.** Rate constants of Mg-ATP-actin monomer association and dissociation at both ends of the actin filament in the absence and presence of side-binding proteins.

| | End | $k_{\text{on}}$ (sub$\cdot\mu\text{M}^{-1}\cdot\text{s}^{-1}$) | $k_{\text{off}}$ (sub$\cdot\text{s}^{-1}$)<sup>†</sup> | $k_{\text{off}}/k_{\text{on}}$ ($\mu\text{M}$) | Reference |
| --- | --- | --- | --- | --- | --- |
| actin alone | Barbed | $11.6 \pm 1.2$ | $1.4 \pm 0.8$ | $0.12 \pm 0.07$ | (Pollard, 1986) |
| | Pointed | $1.3 \pm 0.2$ | $0.8 \pm 0.3$ | $0.6 \pm 0.17$ | (Pollard, 1986) |
| | Barbed | $9.7 \pm 2^*$ | $1 \pm 0.3$ | $0.1 \pm 0.04$ | this work |
| | Pointed | $2.1 \pm 0.8$ | $0.8 \pm 0.4$ | $0.4 \pm 0.35$ | this work |
| VASP | Barbed | $120 \pm 30$ | $1 \pm 3$ | $0.01 \pm 0.03$ | this work |
| | Pointed | $48 \pm 10$ | $0.5 \pm 2$ | $0.01 \pm 0.05$ | this work |
| filamin | Barbed | $8.5 \pm 1.3$ | $0.1 \pm 0.4$ | $0.012 \pm 0.002$ | this work |
| | Pointed | $5.3 \pm 0.1$ | $2.6 \pm 0.2$ | $0.5 \pm 0.04$ | this work |
| α-actinin | Barbed | $7.7 \pm 1.5$ | $0.7 \pm 1$ | $0.1 \pm 0.2$ | this work |
| | Pointed | $0.9 \pm 0.3$ | $0.9 \pm 0.3$ | $1 \pm 1$ | this work |
| NEM-myosin | Barbed | $11 \pm 1$ | $1.6 \pm 0.7$ | $0.15 \pm 0.03$ | this work |
| | Pointed | $0.8 \pm 0.1$ | $0.4 \pm 0.1$ | $0.5 \pm 0.2$ | this work |

<sup>†</sup> all reported dissociation constants from this work are inferred from extrapolation of the elongation velocity as a function of actin concentration to zero concentration, data from Figs. 1, 2 and 4.
<sup>\*</sup> all reported errors from this work are 95% confidence intervals whereas those of (Pollard, 1986) represent SD.

### Kinetic modulation at the pointed-end

Pointed-end association and dissociation rates were estimated in the same manner as those for the barbed-end (Figure 2a). Both the estimated association rates and dissociation rates varied according to the associated side-binding protein used as a tether (Figure 2a and Table 1). The presence of filamin increased the 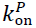 by a factor of ∼5 (from 0.8 in the presence of NEM-myosin to 2.8 sub·μM^−1^·s^−1^). The 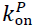 for α-actinin was 0.9 sub·μM^−1^·s^−1^, while, when using VASP, the rate was 44 sub·μM^−1^·s^−1^. On the other hand, the presence of filamin also increased the inferred 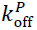 around an order of magnitude from 0.4 (in the presence of NEM-myosin) to 2.6 s^−1^. The inferred 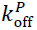 were 0.7 s^−1^ and 8 s^−1^ with α-actinin and VASP respectively.

**Figure 2.**
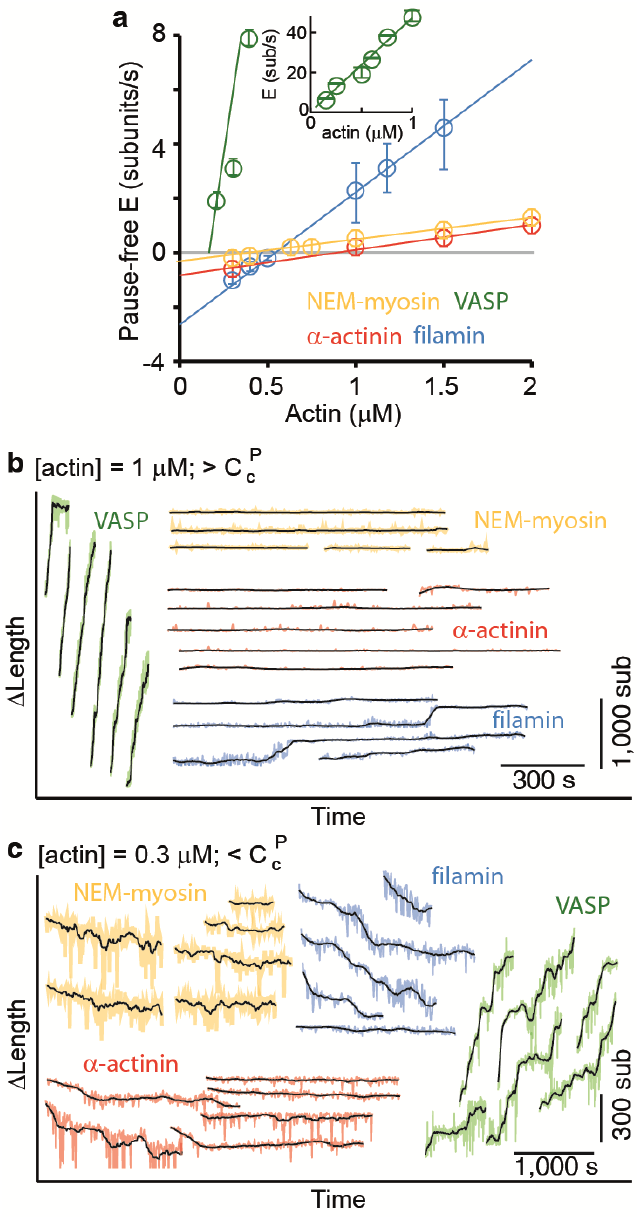
Pointed-end elongation and depolymerization kinetics as a function of the associated side-binding protein. **a**, The elongation velocity (E) is plotted as a function of free actin concentration. Error bars are s.e.m. (n >20). **b-c,** A gallery of traces of Δ*L* as a function of time for pointed-ends observed at (**b**) 1 μM or (**c**) 0.3 μM free actin monomer concentration for the different tethering proteins studied. The raw data are shown in color and the black solid lines are a running average of 10 data points.

Unlike the barbed-end where there were occasional pauses (Figure 1c), the pointed-end displayed mostly a kinetically inactive phase or paused state and only grew sporadically (Figure 2b and c). Such kinetically inactive phases were observed for all free actin concentrations tested. Above the pointed-end critical concentration (e.g. using a free actin concentration of 1 μM), we observed a discontinuous (i.e. growth-pause) behavior for all side-binding proteins (Figure 2b). In the presence of VASP or filamin, pointed-end elongation was readily observed. Pointed-end elongation was much more difficult to visualize when using NEM-myosin and α-actinin (Figure 2b) where elongation occurred for brief periods of time and with slower rates. The elongation velocity during kinetically active phases was influenced strongly by the different tethering proteins used (Figure 2a). Elongation velocity followed the order of VASP >> filamin > α-actinin > NEM-myosin (Figure 2a and b). On the other hand, at 300 nM free actin monomer concentration, i.e. below the pointed-end critical concentration of ∼600 nM, we observed barbed-end growth (Figure 1d) and pointed-end depolymerization (Figure 2c), i.e. treadmilling, in the presence of filamin as a tethering protein (Figure 2c). Treadmilling was also present using NEM-myosin and α-actinin, albeit with slower rates, since pointed-end depolymerization establishes the overall treadmilling rate. Surprisingly, in the presence of VASP, there was no shrinking but polymerization at the pointed-end (Figure 2c). These results suggest that side-binding proteins can also determine actin filament pointed-end growth and depolymerization dynamics.

### The elongation rate varies with occupancy of side-binding proteins

Next, we wanted to study how sensitive filament dynamics are to the presence of each of the proteins tested. Therefore, we measured the elongation rates and pausing as a function of the side-binding protein surface density (Figure 3). For this, we varied the total protein concentration that was allowed to adsorb to the glass surface, therefore changing the number of tethering proteins that interact with a single filament (see Material and Methods and (Crevenna et al., 2008; Howard et al., 1989)). Estimated densities ranged from ∼5 up to ∼20,000 molecules·μm^−2^. At low filamin tether densities (5-200 μm^−2^), elongating actin filaments (at a free actin monomer concentration of 1 μM) showed mostly kinetically active phases (Figure 3a) and an elongation velocity distribution centered at 8.5 sub·s^−1^ (Figure 3a and c). Similar elongation velocity distributions were observed for all tethering proteins tested (Figure 3e and S2-4). At high tether densities (600-20,000 μm^−2^), each side-binding protein tested generated a particular elongation behavior (Figure 3e and Figure S2-4). Using filamin, increasing the surface tether density decreased the mean elongation velocity of kinetically active phases (Figure 3b and d) and increased the fraction of time the filament spent in a paused state, i.e. the pausing probability ‘*P*_p_’ (Figure 3b, d and f). Increasing the VASP density, on the other hand, increased both the elongation velocity and the *P*_p_ (Figure S4). Higher surface concentrations of α-actinin or NEM-myosin had also an effect in elongation velocity (Figure S2-3) and, in addition, NEM-myosin had a strong effect in the *P*_p_ (Figure 3f and Figure S2). These results suggest that a variety of elongation kinetics can arise from the specific interaction of actin filaments with the particular associated side-binding protein.

**Figure 3.**
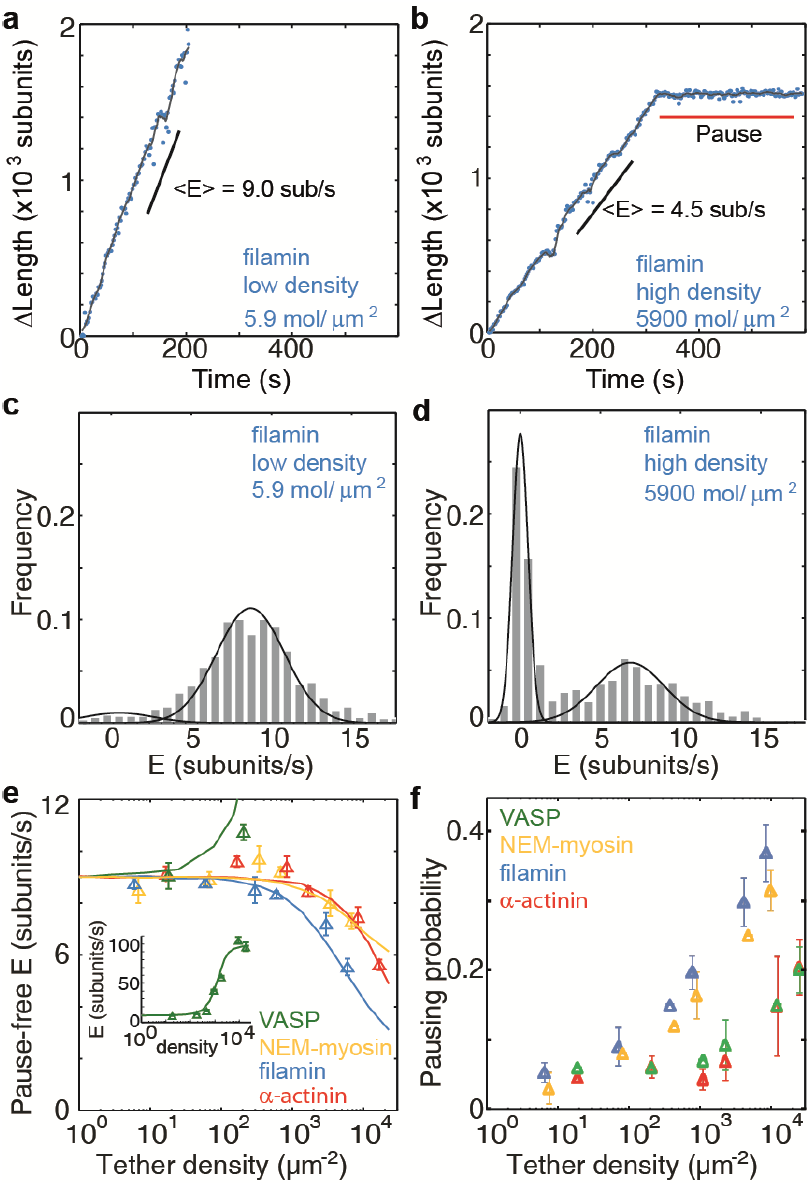
Barbed-end actin filament elongation as a function of the surface density of side-binding proteins. **a-b**. The change in length, Δ*L*, of actin filaments as a function of time for filamin as the surface tethering protein at the (**a**) lowest (5.9 molecules/μm^2^) or (**b**) highest (5900 molecules/μm^2^) density. **c**-**d**, Elongation velocity distribution of filaments using a filamin-coated surface at the (**c**) lowest or (**d**) highest density. Solid lines are fits to Gaussian distributions. The distribution is calculated by binning (0.75 sub/s bin size) the instantaneous elongation velocity of more than 20 filaments. **e**. Elongation velocity as a function of tether surface density, estimated from the kinetically active phases. Solid lines are fits to a model where protein binding induces an allosteric effect that persists along the filament over a certain length scale (see Material and Methods for details). **f**. Pausing probability as a function of surface tether density. Error bars represent s.e.m. (n >20).

### Intrinsic filament dynamics

To verify that the observed kinetic changes and pauses originate from the particular side-binding protein used as a tether, we investigated the intrinsic properties of filament elongation and controlled for artifacts. Single elongating filaments were measured at the lowest protein surface density possible that still allowed filament visualization. At the lowest α-actinin tether density used (5 molecules/μm^−2^, which corresponds to 1 tether molecule every 5-10 microns along the filament), the ends swiveled around their tethering site due to Brownian motion and were clearly free of the surface (Figure 4a). Under these conditions, Barbed-ends showed continuous elongation (Figure 4b) while pointed-ends displayed mostly kinetically inactive state (only 2 of 50 filaments showed growth or depolymerization, Figure 4c-d). For the other tethering proteins, only the paused state was observed on freely swiveling pointed-ends (Figure S5). Using only the pause-free elongation velocities for each actin concentration tested, we estimated association and dissociation rates (slopes in Figure 4e, Table 1). Our estimated values are in good agreement with those previously reported (Table 1). Moreover, the pausing probability, *P*_p_, at either end was insensitive to the actin concentration used (Figure 4f).

**Figure 4.**
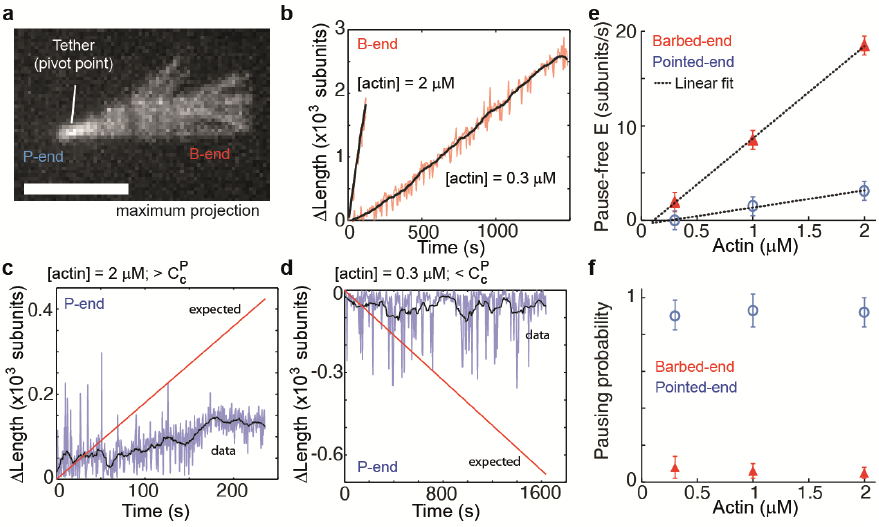
Intrinsic filament dynamics. **a**, A maximum projection image from a movie of an actin filament tethered to a glass surface via a single α-actinin molecule where the tethering position about which the filament swivels is visible as a constriction point. Scale bar: 3 μm. **b**, Change in length of the barbed-end vs. time for individual actin filaments attached to the surface using the lowest tethering protein surface densities at 300 nM and 2 μM concentrations of free actin monomers. **c-d**, Change in length vs. time of elongating single actin filament pointed-ends using the lowest tethering protein surface densities at 2 μM (**c**) or 300 nM (**d**) of free actin monomers. Red lines are the expected elongation based on previously reported rates using NEM-myosin as a tether (Fujiwara et al., 2007; Kuhn and Pollard, 2005). **e**, The pause-free elongation velocity (E) plotted as a function of free actin concentration. The lines represent linear fits. Estimated rates are reported in Table 1. Error bars are s.e.m. (n >20). **f**. Pausing probability as a function of free actin concentration. Error bars represent s.e.m. (n >20).

The low density used for these experiments and the observed pauses on freely swiveling actin filaments (pointed-end only) rules out surface effects (Kuhn and Pollard, 2005) as the determining cause for the pauses at the ends. Another possible source of pauses is light-induced photo-dimerization. From the work of (Niedermayer et al., 2012), it is possible to quantitatively predict the accumulated fraction of filaments where depolymerization has been paused as a consequence of exposure to light (Figure S6). In contrast with this prediction, we observed all swiveling filament pointed-ends, under depolymerizing conditions, to be in a kinetically inactive state at the beginning of image acquisition (N = 40, Figure S6). Only in the presence of a medium to high density of tethering proteins are filament pointed-ends observed to depolymerize (12 of 55, Figure S6).

To further rule out any tether, surface or light-induced effect of the pausing, we used a two-color solution assay to investigate pointed-end growth. Here, a small seed (formed with atto565 labeled actin) was allowed to grow in solution for 15 min in the presence of atto488 labeled monomers, followed by stabilization, dilution and visualization (Figure S7). At a free actin concentration of 1 μM, the concentration used in solution to allow filament elongation, all pointed-ends are expected to grow (at an average rate of ∼0.5 sub/s (Pollard, 1986)). In contrast to this expectation, we observed that only 20% of the seeds grew at the pointed-end (N = 1000, Figure S7). This percentage is higher than we observe in the surface based experiments, which could be due to annealing of filaments in solution (Andrianantoandro et al., 2001; Sept et al., 1999b) or due to lack of the tethering protein. What is clear is that the non-elongating or paused state is not due to either surface or light-induced effects. Taken together, these results show that: a single rate constant describes filament elongation kinetics from ATP-monomers in the absence of side-binding proteins; and the pointed-end has an intrinsic kinetically inactive state.

### Non-kinetic effects of side-binding proteins on filaments

During the course of filament elongation analysis as a function of side-binding protein density on the surface (Figure 4), we noticed that filaments appeared more bent as the tether density increased. To quantify this curviness, we estimated an apparent persistence length 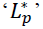 of individual filaments associated with different side-binding proteins (see Extended Supplemental Procedures for details). The persistence length *L*_p_ (Boal, 2012) reflects the material properties of the filament, which are related to its structure (Chu and Voth, 2005, 2006; Pfaendtner et al., 2010), and has already been shown to be tunable by side-binding proteins (such as myosin or cofilin (Bengtsson et al., 2013; McCullough et al., 2008; Murrell and Gardel, 2012)). At the lowest side-binding protein density (∼10 molecules·μm^2^), actin filaments had an 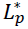 of ∼18 μm and was independent of the associated protein (Figure 5a). At the highest densities (∼10,000 molecules·μm^2^), the presence of NEM-myosin decreased the 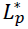 to 4 ± 1 μm while it was reduced to 3 ± 1 μm, 5 ± 1 μm, and 2.2 ± 0.3 μm when using filamin, VASP and α-actinin respectively (N > 50 for each condition, Figure 5a). Moreover, two other phenomena were observed in the presence of side-binding proteins at high density. First, the presence of filamin increased the spontaneous fragmentation of filaments (20 out of 197 filaments vs. less than 1 fragmentation even per 200 filaments) (Figure 5b). Second, barbed-end elongation when tethered with α-actinin had a preference to grow in a counter clockwise direction (Figure 5c). This counter clockwise elongation occurred irrespective of the length the filament had when it landed on the surface. These observations suggest an influence of the tethering protein on the structural properties of the filament.

**Figure 5.**
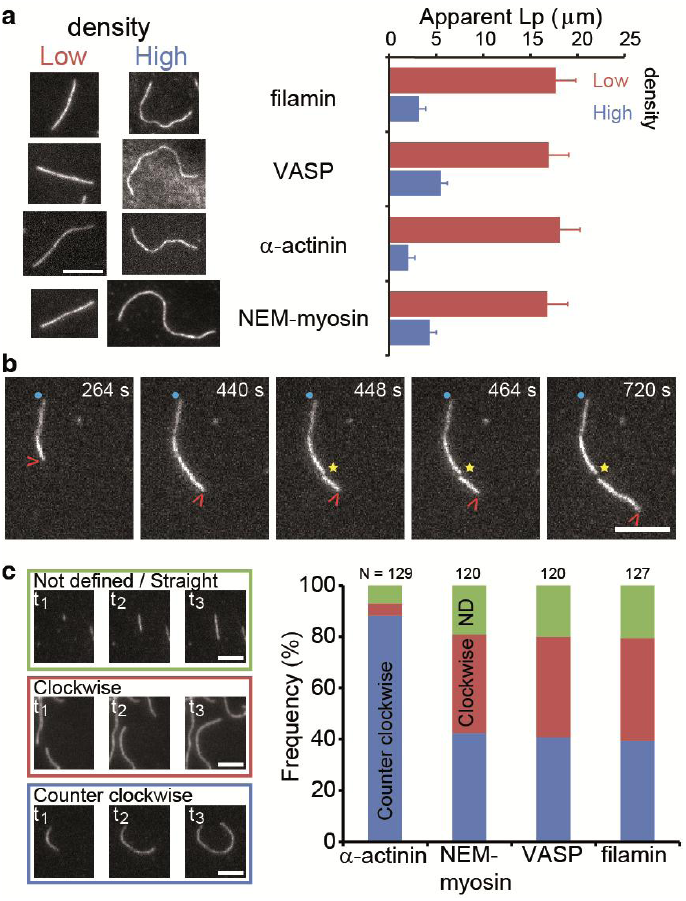
Side-binding proteins alter filament structure. **a**, (left panels) Images of individual filaments attached to the surface using different side-binding proteins at the lowest or highest surface density of tethering protein. Scale bar: 5 μm. (right panels) Estimated apparent persistence length from the angular correlation along the filament contour length at the lowest and highest lattice-binding protein densities. Error bars represent s.e.m. of more than 50 filaments measured per experimental condition. **b**, Images from a movie of an individual growing actin filament under treadmilling conditions. The barbed-end is marked with a red arrowhead and pointed-end with a blue dot. The filament undergoes a fragmentation event (yellow star) at 488 s depolymerizes from its new pointed-end after that while the newly created barbed-end does not elongate. Here, actin concentration was 400 nM. Time is given in seconds. Scale bar: 5 μm. **c,** The direction of filament growth depends on the tethering protein. Barbed-end filament growth direction was classified as straight/not defined, clockwise or counter clockwise from experiments at the highest surface tether density. Examples of each class are shown in the right panels. Scale bars: 3 μm.

Changes in the structure of the filament by binding proteins are known to be able to propagate over several subunits (Orlova et al., 1995). We hypothesize that structural alteration might be the origin of kinetic modulation. To test how well this interpretation could explain our results, we constructed a simple model based on long-range structural alterations to describe elongation as a function of tether density (Figure 3e). We assumed that the structural alteration upon binding propagates linearly along the filament over a certain distance and that this structural effect gives rise to a kinetic change at the ends (Figure S8 and Materials and Methods for details). The model satisfactorily accounts for our experimental results (Figure 3e solid lines) and provides an estimate for the propagation length. Alternatively, we also considered the possibility that VASP acts via a ‘local increase in monomer concentration’ model similar to (Breitsprecher et al., 2011) (see Materials and Methods for details). This local concentration model did not account for the tether density dependence of elongation velocity (Figure S9). Our estimate of the propagation length is a lower limit since considering additional factors in our assay such as tether unbinding and alternative tether density calculations (also see Materials and Methods for details) will only increase the measured propagation length. The induced effects are local-to-short-ranged (∼2-11 monomers) for α-actinin, filamin and NEM-myosin while they are long-ranged (160 monomers) when using VASP (Table S1). Collectively, these results suggest that the side-binding proteins tested alter the structure of the filament.

### Kinetic variability and structural diversity

Next, we wanted to explore whether different conformations could originate diverse kinetics. To gain insight into the plausible range of kinetics an actin filament can exhibit, we performed Brownian Dynamics computer simulations using the different filament models available (Figure 6a). Brownian Dynamics simulations have suggested an electrostatic contribution to the filament asymmetry (Sept et al., 1999a) as well as mapped the kinetics and thermodynamics of filament nucleation (Sept and McCammon, 2001). For each filament model, we calculated the diffusion-limited association rate using a single actin monomer structure in the presence of ATP (see the Material and Methods for details and Table S2 for filament and monomer models used). Unfortunately, given differences in atom pairs used to calculate the association rates, our results were not comparable to those reported previously (Sept et al., 1999a) (see Materials and Methods for details and Figure S10). The association rate at both ends was found to vary over an order-of-magnitude for the different filament structures (Figure 6b). Overall, some models predicted qualitatively well association at the barbed-end, pointed-end or an asymmetry ratio 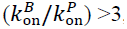, consistent with experiments. Of all filament models, only the Namba and the mode 3 were the closest to agree quantitatively with our data and reported rates (Figure 6b and Table 1). The filament structure has an effect on the elongation kinetics predicted from it.

**Figure 6.**
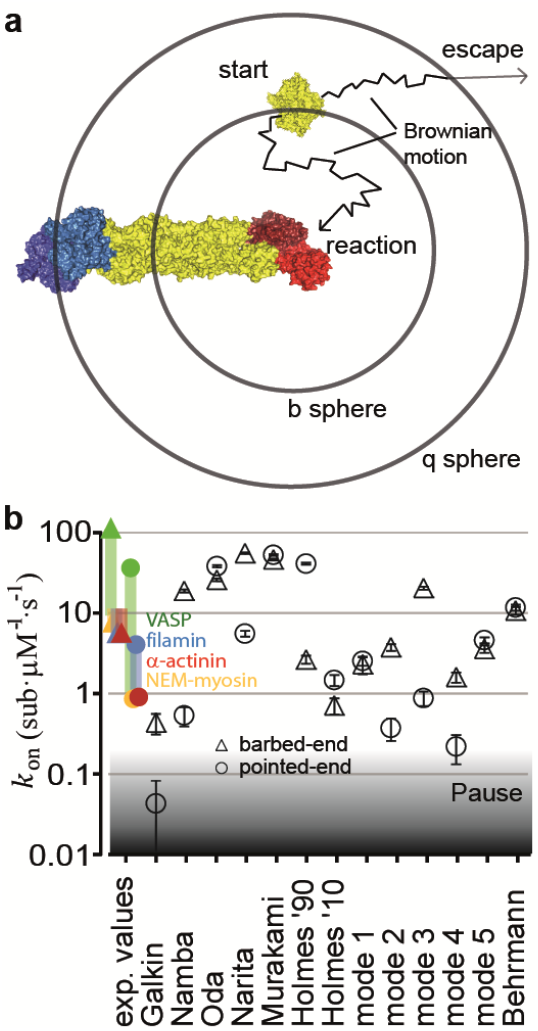
Various filament structures give rise to different kinetics at both ends. **a**, Schematic of the Brownian dynamics computer simulations used to calculate the association rates of an actin monomer to a filament end (see Material and Methods for details). **b**, The calculated association kinetics of the barbed-end (red triangles) and pointed-end (blue circles) determined from the different filament models are shown. For comparison, the experimentally measured values at high tether density are shown as symbols and the range of elongation rates determined from the different surface densities is plotted as a thick line). Rates below 2×10^5^ sub·μM^−1^·s^−1^ become progressively harder to distinguish from paused states when using TIRF microscopy. Error bars represent the upper and lower estimates from the calculations.

Of the filament models that predict the correct asymmetry, most contain monomers which their DNAase binding loop ‘DB loop’ in subdomain 2 forms an α-helix. This result suggests that the DB loop may also play a role in actin filament elongation.

There were combinations of actin monomer structures and filament models which predict association rates at or below ∼0.2 sub·μM^−1^·s^−1^ (Figure 6b and Figure S11), which would appear as paused states when imaged by TIRF microscopy. This result would suggest that the filament structure could be the underlying factor behind elongation pauses.

The variability in predicted association rates as a function of filament model and rates below ∼0.2 sub·μM^−1^·s^−1^ were independent of the monomer structure as similar results to those of Figure 6b were obtained using ADP-actin monomer structures (Figure S11a) or an actin monomer derived from either i) a profilin-actin complex (Figure S11b), ii) from the filament model itself (e.g. a monomer from the Murakami filament model) (Figure S11c) or iii) a myosin-actin complex (Figure S11d). The results from these simulations suggest that the lattice structure can determine association rates at the ends of the filament.

## Discussion

Our results provide insights into the origin of the filament asymmetry, a consequence of a non-elongating state at the pointed-end, and on general versatility that actin dynamics respond to binding proteins.

Through accurate measurements of pointed-end association kinetics we have observed that in the absence of tethering proteins and in the presence of VASP yielded equivalent critical concentrations for both ends (∼0.2 μM, Table 1), which implies that detailed balance at equilibrium, i.e. 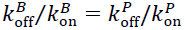 (Hill, 1987), is fulfilled. Therefore, intrinsic elongation or in the presence of immobilized VASP can be thought of as ATP-actin binding to an ATP-actin filament. The existing discrepancy in estimated critical concentrations at both ends (Fujiwara et al., 2007) originates from the presence of a previously uncharacterized kinetically inactive or non-elongating state at the pointed-end. This kinetically inactive state is consistent with a non-elongating structural conformation observed by cryo-electron microscopy (Narita et al., 2011). The kinetic asymmetry of the pure actin filament may be low (∼3) and such non-elongating or closed conformation at the pointed-end would reinforce the effective filament asymmetry. The transition at the pointed-end from the open to the closed state may be coupled to ATP hydrolysis or phosphate release at the terminal subunit. The presence of this open-to-close transition at the pointed-end would explain why the terminal subunit has an estimated different rate of phosphate release compared to the filament lattice (Fujiwara et al., 2007).

Actin filaments in association with any of the four proteins tested displayed a change in elongation velocity, an increase in pausing and a change in filament flexibility. Therefore, it is possible that these three characteristics have a common origin. For three of these filament-binding proteins (myosin, α-actinin and filamin), the binding interface to the actin filament is formed by two consecutive monomers along the same strand (Galkin et al., 2008; Galkin et al., 2010a; Lorenz and Holmes, 2010). These side-binding proteins might directly occlude the binding site for the next monomer either partially (reducing the elongation velocity) or completely (giving rise to an elongation pause). A partial filament distortion could become into a defect that propagates in the lattice decreasing the observed filament stiffness and may impact the association rate. In this respect, side-binding proteins could be thought of as allosteric regulators of actin filament kinetics. In this respect, actin filaments are known to be subject to allosteric regulation by other associated proteins (Egelman and Orlova, 1995; Galkin et al., 2012). In particular, myosin (Prochniewicz and Thomas, 1997), cofilin (Galkin et al., 2001; Prochniewicz et al., 2005), dystrophin (Prochniewicz et al., 2009) and utrophin (Prochniewicz et al., 2009) are known to induce structural changes to the actin filament. Similar to filamin and α-actinin, dystrophin and utrophin bind actin through calponin-homology (CH) domains (Galkin et al., 2010a). Moreover, binding to the filament is cooperative for cofilin (De La Cruz, 2005), αE-catenin (Hansen et al., 2013) and myosin (Orlova and Egelman, 1997). The basis for this allosteric regulation could originate from the stabilization of an existing structural state of the filament (Galkin et al., 2001), given that the actin filament is structurally polymorphic (Galkin et al., 2010b). Therefore, it is possible that the observed elongation kinetics and pauses arise from direct modulation of the filament structure. In line with this hypothesis, two other proteins, the actin-binding domain of αE-catenin (Hansen et al., 2013) and an N-WASP construct (Khanduja and Kuhn, 2014), have recently been shown to alter filament kinetics and one of them, the actin-binding domain of αE-catenin, also influences filament structure (Hansen et al., 2013). Although more and more atomically accurate simulations are required to understand the molecular basis of monomer association and dissociation from the filament ends, our results provide evidence that lattice structural changes can have strong effects in actin filament growth kinetics.

Our experimental approach, that of tethers immobilized on a solid surface, imposes geometric constrains on filament growth that may not faithfully represent the cytosol. However, actin filaments form part of the cell cortex (ref) or of focal adhesions (Kanchanawong et al., 2010) where they assemble into an oligomeric membrane-anchored complexes with many actin-binding proteins tethered to the plasma membrane surface. Moreover, the cell interior is very crowded (Luby-Phelps, 2000), and some sub-cellular actin arrays are tightly packed (Jasnin et al., 2013) both conditions which may immobilize actin-binding proteins and generate similar constrains during filament growth. Depending on the local cross-linker protein abundance in the cell, turnover kinetics on the order of 1 μm of filament within ∼1 minute can be achieved, a rate at which treadmilling could become a contributing factor to cellular retrograde flow in the lamellipodium (Ponti et al., 2004; Watanabe and Mitchison, 2002). Additionally, filament structural changes generated by side-binding proteins may also play a more active role in the identity and turnover of actin-based sub-cellular structures than previously thought, by regulating processes such as branching and fragmentation (Hansen et al., 2013) or network mechanics (Jensen et al., 2014). Given the vast number of side-binding proteins, kinetic modulation via structural alteration may be a general regulatory mechanism of actin dynamics.

## Materials and Methods

### Proteins

Actin was obtained from acetone powder, made from rabbit skeletal muscle, by one round of polymerization and pelleting by centrifugation (Spudich and Watt, 1971). The resulting pellet was depolymerized in G-buffer (1 mM Tris-HCl pH 7.8, 2 mM ATP, 2 mM CaCl_2_, 2 mM DTT) overnight at 4 °C followed by gel filtration on a Sephacryl S-300 column. Myosin was purified and chemically inactivated with N-Ethyl-Maleimide according to the published protocol (Breitsprecher et al., 2009b). Atto488-actin, α-actinin, and filamin were purchased from Hypermol (Bielefeld, Germany). Alternatively, actin was labeled with succinimidyl ester atto488 (ATTO-TEC GmbH, Germany) on random lysine residues. Actin labeling was performed under polymerization conditions (50 mM KCl and 2 mM MgCl_2_) followed by depolymerization and gel filtration in G-buffer. Unlabeled and labeled actin were mixed to yield a final ratio of 2:1 unlabeled:labeled actin. The actin mixture (20 μL) was snap frozen and stored at −80 °C until further use. Before use, an actin aliquot was centrifuged to remove possible aggregates.

A plasmid containing the gene of *Dd* VASP was kindly provided by J. Faix, (Hanover, Germany). VASP was expressed using a pCoofy plasmid in Sf9 cells with a MBP-tag, and purified following standard methods as described previously (Scholz et al., 2013). MBP-VASP was used without cleavage, since removal of the tag resulted in protein aggregation and degradation.

### Imaging

Flow cells were made as a sandwich of a cover slip (20 × 20 mm), parafilm with an approximately 5 mm wide channel and a glass slide. A. The surfaces of the flow cells were passivated to avoid adsorption of actin to the sample holder by incubating them with 10% (w/v) of BSA in PBS for 10 min. Flow cells were washed three times with 90 μL of G-buffer. The tethering protein was then applied for 5 min and the flow cell was then washed again three times with 90 μL of G-buffer. Actin (33% atto488-actin) was incubated 5 min on ice with 1/10 volume of 10x ME buffer (400 μM MgCl_2_ and 2 mM EGTA) to exchange Ca^2+^ for Mg^2+^. The actin-containing solution was mixed with imaging buffer (catalase, β-mercapto ethanol, glucose oxidase, 0.8% (v/v) D-glucose, 0.25% (w/v) methylcellulose, and 1/10 volume of 10x KMEI buffer (500 mM KCl, 20 mM MgCl_2_, 20 mM EGTA and 300 mM imidazole), with a final pH of 7.1) and introduced into the flow cell. TIRF microcopy was performed using a TILL photonics inverted microscope. A single actin aliquot was used within 12 hours.

Lattice-binding protein surface density was estimated from the protein concentration, the sample volume (∼10 μL) and the surface to which the sample was adsorbed (a flow cell of 5 mm × 20 mm, giving 100 mm^2^) as done previously (Crevenna et al., 2008; Howard et al., 1989). All protein in solution was assumed to adsorb on the upper and lower glass surfaces To achieve consecutive lower tether densities, the total protein concentration was serially diluted. At low tethering protein concentration, individual filaments swiveled around distinctive attachment points indicating that they are bound to single tethering molecules as observed previously (Crevenna et al., 2008; Howard et al., 1989). To estimate the density in an alternative manner, we measured the average number of pivot points per micron of filament at the two lowest protein concentrations and divided that by the average area covered during swiveling. Assuming a linear scaling with protein concentration, this estimation resulted in a slightly lower density (by a factor of 2) compared to those reported in Fig. 3. As the concentration-based estimated densities represent an upper limit and are easy to reproduce, we report those in the text.

### Two color solution assay

Filaments were formed using atto565-labeled actin in G-buffer by addition of 1/10 volume of 10x KMEI buffer. After >2h of polymerization at room temperature, filaments were fragmented by shearing and subsequently mixed with atto488-labeled monomers and allowed to elongate for 15 min. Filaments were then stabilized with unlabeled phalloidin and diluted for imaging on an Epi-Fluorescent Microscope (Axiovert 200, Zeiss).

### Data analysis

Raw movies were corrected for *x*- and *y*-stage drift by first calculating its magnitude via image correlation spectroscopy (Hebert et al., 2005) and, secondly, correcting the drift by bicubic interpolation. Drift estimation and correction were implemented in custom programs written in LabView and MATLAB. Kymographs of single filaments were made using Metamorph or Image J, while further analysis and Monte Carlo simulations were carried out using MATLAB. The position of the filament tip, per line in the kymograph, was estimated by fitting an error function as previously described (Demchouk et al., 2011). More than 20 filaments were analyzed per condition. To estimate the first pause distribution, we used the model described by (Niedermayer et al., 2012) with *ω* = 2 × 10^6^. The light intensity for treadmilling experiments ranged from 0.74 to 0.92 mW·mm^−2^. Growth orientation was assessed manually.

### Brownian Dynamics simulations

In order to compute the association rates, Brownian dynamics (BD) simulations of different filament models were performed using BrownDye software (Huber and McCammon, 2010). The filament models were built using the coordinates of the following PDB codes: 3G37 (Murakami), 2Y83 (Narita), 3J0S (Galkin), 2ZWH (Oda), 3MFP (Namba), 1ATN (Holmes ‘90), 1J6Z (holmes ‘10), 4A7N (Behrmann) and modes 1-5 from Ref (Galkin et al., 2010b). See Table S2 for details. Any molecules other than actin or nucleotide (ATP, ADP) were deleted from the structures in order to have fully comparable models. In addition, differences in the sequence of the N-terminus of chicken skeletal muscle actin (for the Galkin model: PDB 3J0S) were converted to that of rabbit skeletal muscle actin (PDB:1ATN) by making the following substitutions: D2E, D4E, I5T and A6T. PQR files were generated using the software pdb2qpr (Dolinsky et al., 2004). Electrostatic potentials were computed by solving the nonlinear Poisson-Boltzmann equation with APBS version 1.4 (Baker et al., 2001) using the Amber force field at 298 K. ATP and ADP partial charges were taken from the internet (http://www.pharmacy.manchester.ac.uk/bryce/amber). Atomic charges were mapped to grid points via cubic B-spline discretization. The relative permittivities used for the solute and solvent were 4 and 78.5, respectively. All the electrostatic potential calculations were performed assuming 50 mM KCl. Between 1 and 3 million trajectories were performed for the calculation of each association rate. A single trajectory was considered successful (i.e. the complex was formed) when at least 3 polar contact pairs reached a distance of ≤ 10 Å (Sept et al., 1999a). The list of polar contact pairs was derived from the input coordinates of each filament model. A polar contact pair is defined as a pair of atoms within 3.5 Å from each other where one atom is from the monomer and the other is from the filament. The complete lists of contact pairs are provided in supplementary Table S3.

Although current simulations still have inaccuracies (e.g. neglect molecular flexibility, contact criteria definition, and approximated solution of electrostatic interactions) that may limit the reliability of the results, previous Brownian Dynamics simulations have been successful in describing experimental data for the association of actin to actin binding proteins (Pang et al., 2012) as well as for several other complexes (Gabdoulline and Wade, 2001).

The models built in ’90 and ’10 by Holmes contained steric clashes between residues P243 and T203 of the *n^th^* monomer element of the filament and residues around P322 and D288 of the *n* + 2 monomer element (Figure S8). We removed these clashes by running 300 steps using energy minimization steepest descendent in GROMACS 4.5 (Hess et al., 2008; Pronk et al., 2013) using the Amber99sb force field (Figure S8).

Unfortunately, it was not possible to compare our calculated rates with previous calculations (Sept et al., 1999a) due to discrepancies in the atom pairs used to define a contact within the bound structure. As an example, the pairs P243O-K291NZ and V46O-K291NZ were reported in (Sept et al., 1999a) to be separated by 2.8Å and 3.7Å respectively, whereas we found them to be 10.6Å and 16.6Å respectively (Figure S9).

### Persistence Length calculation

Individual filaments were extracted from the measured data using open active contours with JFilamin, a plug in for Image J (Li et al.; Smith et al.). The filament persistence length ‘*L*_p_’ was determined by calculating the angular correlation (Isambert et al., 1995):

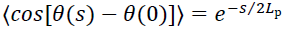

where the brackets represent the average correlation function of the tangent *θ*, measured along the contour length *s*. The point spacing used to reconstruct a single filament was between 6 and 10 points per micron to avoid artifacts in the *L*_p_ estimation (Isambert et al., 1995; McCullough et al., 2008; Smith et al., 2010). All data analysis was done in MATLAB.

### Models for describing the elongation rate as a function of side-binding protein density

To account for the experimentally observed elongation rates as a function of the density of tethering proteins (Figure 3e), we tested different models.

#### Local change in actin concentration

First, the strong increase in the elongation rate in presence of e.g. VASP as a tethering protein was thought to be related to the multiple actin monomer binding sites on each VASP protein (Breitsprecher et al.; Breitsprecher et al.; Hansen and Mullins). Theoretically, the polymerization kinetics is expected to be inhomogeneous along the actin filament and only to be locally enhanced through a higher local concentration of free actin monomers due to the presence of VASP. The growth at one end of the polymer can be written as follows

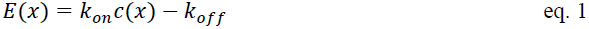

where *E*(*x*) is the elongation rate at the position *x* along the filament axis, *c*(*x*) the local concentration of globular actin at this position and *k_on_* and *k_off_* are the association and dissociation rates. The average elongation rate along the filament length is given by

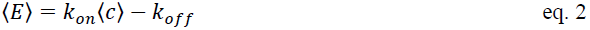

where 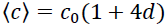, *c_0_* is the free actin concentration in solution and *d* is the density of VASP protein at the surface. Given that one VASP protein can bind 4 actin monomers, the local concentration of actin monomers available for polymerization can be significantly increased at the site of a tethering protein. The relationship between the average elongation rate and the protein surface density was expected to be linear as follows

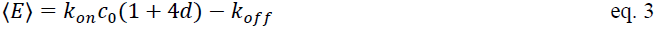

Experimental points do not show a linear dependency on the protein surface density as expected from this model (Figure S11). An alternative mode of operation for VASP has been recently postulated where the protein does not only increase the local concentration but also transfers monomers from its monomer binding domains to the filament tip (Breitsprecher et al., 2011). The surface density dependency of this alternative model would, nonetheless, predict a linear behavior as well albeit with a different slope. In addition, this local increased model would not explain the effect observed with filamin and α-actinin used as tethering proteins, where a decrease in the elongation rate was observed.

#### Propagation of modified association rates induced by binding of side-binding proteins

The following alternative model was proposed to explain enhancement or hindering of elongation. The interaction between the growing filament and a tethering protein was thought to give rise to a modified association rate at the site of interaction. This modified elongation was then allowed to propagate over a certain distance *L_C_* (Figure S12). The decay of the interaction effect was modeled as linear, which would be expected for the release of torsional stress.

The polymerization of actin filaments at the barbed end was simulated using the Monte Carlo method. The average elongation rate was calculated from the length of the polymer over time for each condition. The effect of the tethering protein was simulated by a change in the effective *k*_on_ at the site and vicinity of the tethering protein (Figure S8). First, the tethering protein’s positions were randomly chosen according to the protein density. The surface density was calculated assuming that all added tethering protein adsorbed to the surface and was functionally active. To convert from surface density to fractional occupancy, we used the area occupied by 1 μm of actin filament (0.006 μm^2^), using a value of 370 subunits per micron of filament. Possible dissociation of the filament from the tethers was neglected. During the simulated polymerization, the effective *k*_on_ was changed to 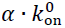 at the position of the tethering protein and decreased linearly until reaching the free actin value of 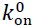 after a characteristic length (*L*_C_) counted in monomers of actin. The values of 11 μM^−1^s^−1^ and 2 s^−1^ for barbed-end 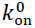 and *k*_off_ respectively were taken from the literature (Pollard, 1986) and the average elongation rate was taken from measurements at the lowest tethering protein density (Figure 3e). The only free parameters were α and *L*_C_, which were determined for each curve (i.e. elongation as a function of lattice-binding protein density) by comparing the simulated elongation rates to the experimental values and minimizing the χ^2^. A first round of simulations were performed to roughly estimate the optimal fit parameters followed by a second set of simulations to obtain a better resolution on α and L_C_. Results obtained are shown in Supplementary Table 1 with 68% confidence bounds. As our simple model most likely overestimates the number of bound side-binding proteins, the estimated characteristic length provides a lower limit for the propagation length of the interaction.

## Acknowledgements

We thank D. Kovar for the Image J macros’s to perform the filament analysis, G. F. Schroeder for sharing pdb files, J. Faix for the DdVASP expression plasmid and J. Dominguez for valuable discussions. This work was supported by the Max Planck Society and by grants from the Deutsche Forschungsgemeinschaft through the SFB 1035 (to OFL) and the SPP 1464 (to RWS and DCL), a CONACYT-DAAD scholarship A/09/7253 (to MA), the Ludwig-Maximilians-Universität München (LMUInnovativBioImaging Network) and the Nanosystems Initative Munich (NIM).

